# The evolutionary dynamics of alternative splicing during primate neuronal differentiation

**DOI:** 10.1101/2024.02.20.581203

**Authors:** Alex Ritter, Andrew Wallace, Neda Ronaghi, Jeremy R Sanford

## Abstract

Alternative splicing (AS) is emerging as an important regulatory process for complex biological processes such as neuronal differentiation. To uncover the functional consequences of AS during neuronal differentiation we performed a comparative transcriptomic analysis using human, rhesus, chimpanzee and orangutan pluripotent stem cells. Transcriptomic studies commonly involve the identification and quantification of alternative processing events, but the need for predicting the functional consequences of changes to the relative inclusion of alternative events remains largely unaddressed. Many tools exist for the former task, albeit often limited to rudimentary event types. Few tools exist for the latter task; each with significant limitations. To address these issues we developed junctionCounts, which captures both simple and complex pairwise AS events and quantifies them with straightforward exon-exon and exon-intron junction reads in RNA-seq data, performing competently among similar tools in terms of sensitivity, false discovery and quantification accuracy. Its partner utility, cdsInsertion identifies transcript coding sequence information, including the presence of premature termination codons, gathered via *in silico* translation from annotated start codons. It then couples transcript-level information to AS events to predict functional effects, i.e. nonsense-mediated decay (NMD). We used junctionCounts and related tools to discover both conserved and species-specific splicing dynamics as well as regulation of NMD during differentiation. Our work demonstrates this tool’s capacity to robustly characterize AS and bridge the gap of predicting its potential effect on mRNA isoform fate.

**GRAPHICAL ABSTRACT:** 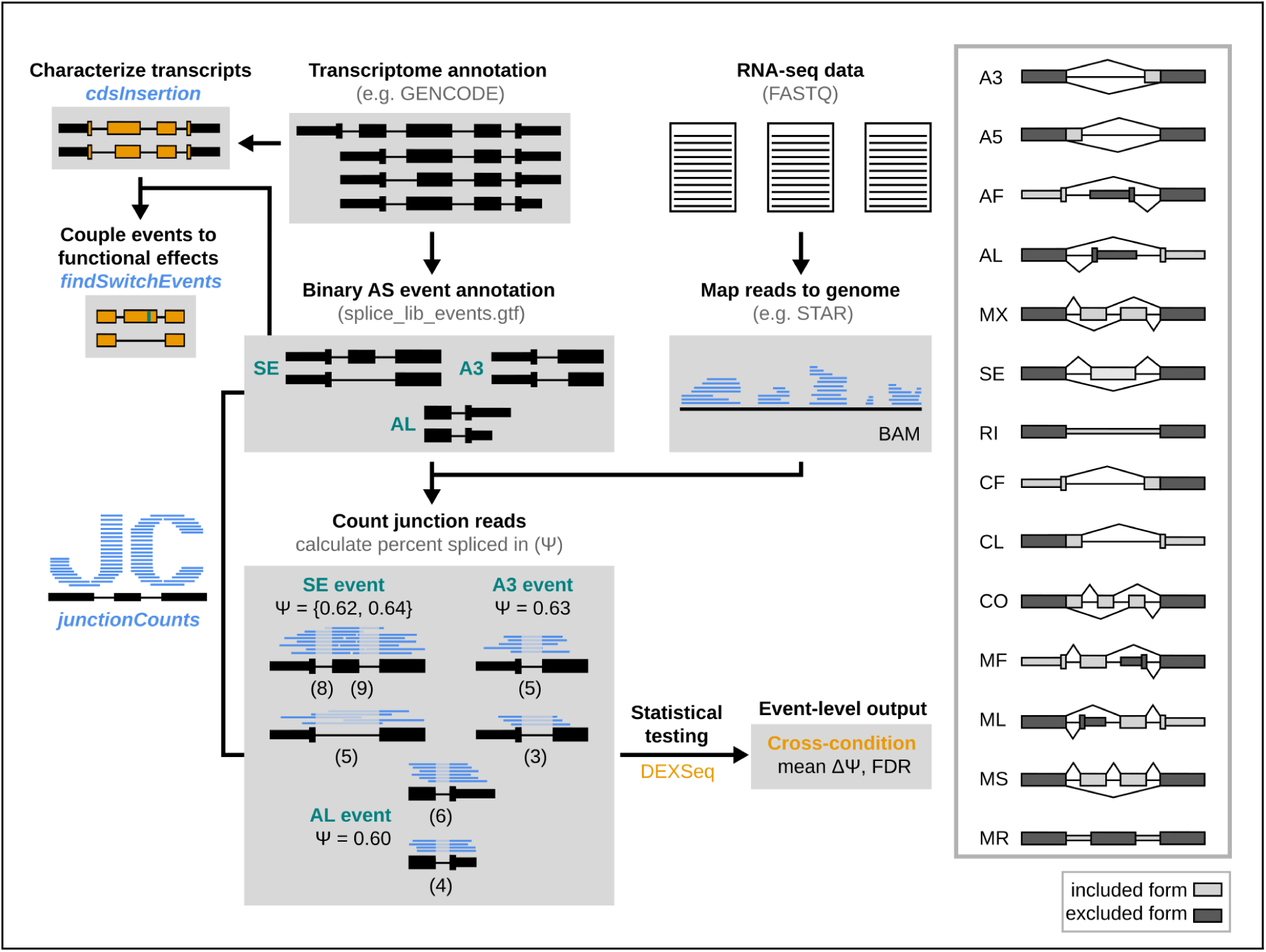

junctionCounts is an alternative splicing analysis tool that identifies both simple and complex splicing events from a gene annotation and then measures their percent spliced-in from mapped RNA-seq junction reads.

## INTRODUCTION

Alternative splicing (AS) generates a diverse array of mRNA isoforms from a single locus. The consequences of this process can have manifold effects on gene expression by altering mRNA half-life (1), intracellular localization (2), translation efficiency (3), and most obviously, by producing different protein isoforms (4). AS plays critical roles in a variety of biological processes including disease pathology, cancer, and cellular differentiation. In cellular homeostasis, AS is tightly regulated to control the precise expression of diverse mRNA isoforms, allowing cells to adapt to changing conditions and to achieve different states of activation in immune cells, for example (5). Regulatory elements, including *cis*-acting splicing enhancers and silencers, as well as *trans*-acting splicing factors, coordinate the inclusion or exclusion of alternative exons or splice sites during transcription (6).

Dysregulation of AS can contribute to the production of aberrant protein isoforms, impacting critical cellular functions. In cancer, this can result in the generation of oncogenic isoforms, altered signaling pathways, and evasion of regulatory mechanisms (7). Often, mutations in splicing factors and in *cis*-elements underlie oncogenic AS dysregulation (8). During cellular differentiation from pluripotent stem cells, AS orchestrates the precise control of gene expression, directing the development of specialized cell types with distinct functions (9). This process is intricately involved in shaping the cellular landscape, driving fate decisions, and maintaining tissue homeostasis (10). The important impact of AS in disease and cellular differentiation underscores its significance as a regulatory force in biological processes and highlights its potential as a therapeutic target in pathological conditions.

It is thus important, in any eukaryotic cellular context, to understand gene expression at the isoform level. The complex nature of AS, however, creates numerous obstacles to its accurate study. mRNA isoforms can, and have canonically been, characterized in terms of binary events that either include or exclude an alternative feature (exon, intron or splice site). This mode of characterization faces the challenge of differentiating between complex and overlapping event features, and also relies on comprehensively annotated gene structures. The latter problem is highlighted by the lack of records for transcripts clearly supported by mapped reads in available references. To address this, reference-guided or *de novo* transcriptome assembly has become a widely used step in RNA-seq analysis. In this process, transcript structure is predicted from the data with or without the use of a reference gene annotation as a template (11). This process allows analysts to consider potential novel transcripts and novel alternative event features that may be important to the biological phenomena under study.

Upon establishing comprehensive gene models, it subsequently becomes important to understand how AS configures the coding and noncoding regions of resultant mRNA isoforms. Unfortunately, common tools for transcriptome assembly (12, 13) are unable to provide information on the presence and nature of open reading frames (ORFs) that may be contained within predicted novel transcripts. One example of an available tool, Transdecoder (14), somewhat addresses this limitation – however, it was developed for use with Trinity (14) which is intended for completely *de novo* transcriptome assembly in the absence of a reference genome assembly or any annotations. Consequently, Transdecoder performs *de novo* ORF prediction with intent towards identifying all ORFs that could convincingly give rise to proteins. In most well-annotated genomes, however, existing ORF annotations are available for the majority of genes in which new transcripts might be identified. In these cases a potentially more reliable approach is to examine novel coding sequences that begin with high-confidence annotated start codons. This prediction approach is useful not only because it informs on potential novel peptides, but also because it has the ability to identify the presence of premature termination codons (PTCs) or high-confidence start codons that lack an in-frame downstream stop codon. In these latter cases, translation is expected to result in surveillance of host transcripts via nonsense-mediated decay (NMD) and non-stop decay (NSD) respectively.

NMD exemplifies an important potential outcome of AS. This translation-dependent surveillance mechanism identifies transcripts containing PTCs ≥ 55 nt upstream of a splice junction and triggers its degradation and translational suppression (15). PTCs can be introduced through AS by inclusion of PTC-containing exons (poison exons), through splicing events that shift the reading frame of the message and by splicing events occurring within the 3’ untranslated region (3’UTR) (16). Other outcomes that can dramatically affect the fate of mRNAs may involve coding-to-noncoding switches, long-to-short UTR switches, or the inclusion of rare codons. It is thus important to profile mRNA coding features, and to connect binary AS events to their potential impacts to mRNA fate and function. The functional impacts of AS, however, remain difficult to predict.

To address this problem we developed junctionCounts comprising: junctionCounts event identification and quantification modules, cdsInsertion and findSwitchEvents. junctionCounts identifies and quantifies a diverse array of AS events. cdsinsertion translates provided transcripts *in silico* from user-provided overlapping start codons and determines resulting transcript characteristics such as UTR lengths, putative primary structure, the presence of PTCs, PTC distances from downstream splice sites, and more. Its partner utility, findSwitchEvents, bridges the gap between transcript-and event-level analysis, allowing one to take advantage of the potential superior quantification accuracy and regulatory interpretation of event-level analysis while still leveraging information that can only be derived from full-length transcripts (17). In this study, we present junctionCounts as a powerful and flexible tool for studying AS in a variety of cellular contexts, and we demonstrate its utility in not only identifying significant splicing events, but inferring their functional outcomes.

## MATERIALS AND METHODS

### Alternative event definition in junctionCounts

Alternative events are defined as instances in which pairs of: a) identical upstream 5’-and identical downstream 3’-exon boundaries, b) non-identical upstream 5’-transcript termini and identical downstream 3’-exon boundaries or c) identical 5’-exon boundaries and non-identical 3’-transcript termini are separated by any two combinations of distinct exon coordinates. Cases in which the aforementioned pairs are separated by more than two sets of distinct exon coordinates result in distinct alternative events for all pairwise combinations of those sets. Cases in which two or more events share the same splicing structure result in a single representative event in which the most proximal outer exon boundaries are used. This approach to event identification encompasses standard alternative event types and further identifies non-standard events of complex exon structure.

### Alternative event classification in junctionCounts

The majority of event types in junctionCounts correspond to types explicitly defined elsewhere (18). junctionCounts also adopts previous usage of the term ”complex” (19, 20) to refer to non-standard event types, distinguishing between internal (CO), 5’-terminal (CF), and 3’-terminal (CL) contexts. Below are full descriptions of the criteria for each event type:

**SE** - *skipped exon*: an event in which a pair of 3’- and 5’-splice sites are separated by a single exon (the skipped exon) in one isoform and spliced directly together in another. The form containing the intermediate exon is the included form.
**MS** - *multiple skipped exons*: an event in which a pair of 3’- and 5’-splice sites are separated by multiple exons in one isoform, and spliced directly together in another. The form containing the intermediate exons is the included form.
**A3** - *alternative 3’-splice site*: an event in which an upstream exon and the 3’-boundary of the downstream exon are common to both isoforms, but the 3’-splice site is distinct. The form containing the most upstream of the alternative 3’-splice sites is the included form.
**A5** - *alternative 5’-splice site*: an event in which a downstream exon and the 5’-boundary of the upstream exon are common to both isoforms, but the 5’-splice site is distinct. The form containing the most downstream of the alternative 5’-splice sites is the included form.
**MX** - *mutually exclusive exons*: an event in which a pair of 3’- and 5’-splice sites are separated by a distinct exon in each isoform. The only requirement for the two exons being distinct is that they do not share either splice site. It is possible for the alternative exons in an MX event to partially and completely overlap, provided their boundaries do not coincide. The form in which the alternative exon’s 3’-splice site is the most upstream of the mutually exclusive exons is the included form.
**RI** - *retained intron*: an event in which a pair of adjacent exons are spliced together in one isoform, but joined together in another by retention of the intron separating them. The form with the retained intron is the included form.
**MR** - *multiple retained intron*: an event in which a set of three or more exons are spliced together in one isoform, but connected in the other by two or more consecutive retained introns. The form with the retained introns is the included form.
**AF** - *alternative first exon*: an event in which each isoform has its own distinct 5’-terminal exon; each with a distinct 5’-terminus and 3’-splice site. The exon immediately downstream of the terminal exon is common to both isoforms. The form with the most upstream 5’-terminus is the included form.
**MF** - *multiple alternative first exons*: an event in which each isoform has its own distinct 5*^’^*-terminal set of one or more exons upstream of a single shared exon. This event type is distinguished from AF in that either isoform must contain more than one unique exon. The form with the most upstream 5’-terminus is the included form.
**AL** - *alternative last exon*: an event in which each isoform has its own distinct 3’-terminal exon (i.e. each with a distinct 3’-terminus and 5’-splice site). The exon immediately upstream of the terminal exons is common to both isoforms. The form with the most downstream 3’-terminus is the included form.
**ML** - *multiple alternative last exons*: an event in which each isoform has its own distinct 3’-terminal set of one or more exons downstream of a single shared exon. This event type is distinguished from AL in that either isoform must contain more than one unique exon. The form with the most downstream 3’-terminus is the included form.
**CO\*** - *complex internal*: this is a general category for events that do not meet any of the above criteria and do not involve transcript termini. The isoform with the longest spliced length is the included form.
**CF\*** - *complex 5’-terminal*: this is a general category for events that do not meet any of the above criteria and involve alternative 5’ transcript termini. The form with the most upstream 5’-terminus is the included form.
**CL\*** - *complex 3’-terminal*: this is a general category for events that do not meet any of the above criteria and involve alternative 3’ transcript termini. The form with the most downstream 3’-terminus is the included form.

**\***Complex event types capture less straightforward cases of binary splicing events that can involve combinations of multiple alternative features (terminal and internal exons and/or splice sites).

### Alternative event quantification in junctionCounts

junctionCounts employs a junction read-centric approach to alternative event quantification. For each read or read pair, junctionCounts considers matches between splice junctions identified in the alignment and event splice junctions, as well as overlaps between contiguous mapped read sequence and informative exon-intron junctions. Informative exon-intron junctions are those that are overlapped by an exon in the alternative isoform. Reads overlapping such an exon-intron junction are considered consistent with the alternative isoform. Key examples of this occur in the excluded isoform of RI events, which are overlapped by the exon of the included form. After establishing the event isoforms with which a read is consistent, junctionCounts attempts to disambiguate the read assignment using exon-exon and exon-intron junctions that are unique to specific isoforms, when possible. With this approach, junctionCounts goes beyond simple junction-by-junction read counting. Both the event and the informative exon-intron junction definition prohibit scenarios in which reads are assigned to both isoforms of the same event. With read-to-event isoform consistencies established, read counts are tallied for each exon-exon and informative exon-intron junction for each isoform of each event. A percent spliced in (PSI or Ψ) value is calculated for all pairwise combinations of included and excluded junction counts, yielding the ratio between an included form’s junction counts and the sum of the included *and* excluded form’s junction counts:

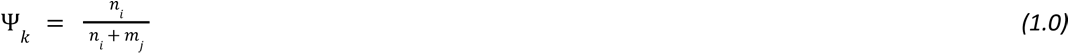

Where *n_i_* is the number of reads assigned to the included form junction *i* and *m_j_* is the number of reads assigned to the excluded form junction *j*. A set of PSI values is established for each sample. Additionally, junctionCounts calculates the minimum (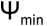), maximum (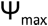) and span PSI (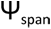):

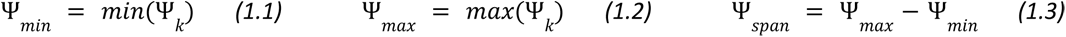

The 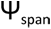 serves as a rough measure of within-sample uncertainty. junctionCounts reports these values as well as the included and excluded junction read counts (*n* and *m* in Equation 1.0) for each event. As an optional method of assessing within-sample uncertainty, junctionCounts offers bootstrap quantification, which repeats a user-specified number of rounds of bootstrap read selection and re-quantification. For each bootstrap round, junctionCounts reports all measurements, in addition to the initial non-resampled quantification. Statistical testing of events between conditions in experiments with at least two replicates per condition is done by comparing the dispersions of included versus excluded form junction counts using DEXSeq (21), yielding per-event changes in PSI (dPSI or 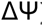) and positive false discovery rate (Q-value).

### Benchmarking junctionCounts event identification and quantification

junctionCounts was evaluated based on its performance relative to two established splicing analysis tools: MAJIQ (22) and splAdder (23). Each of the three tools were used to identify and quantify events in mouse cerebellum and liver RNA-seq data in triplicate from Vaquero-Garcia et al. (2016) (22) who replicated the experiments in Zhang et al. (2014) (24). Corresponding RT-PCR data, from Vaquero-Garcia et al. (2016) (22), for 50 SE events were used to measure the linearity of measured PSI and dPSI values with biologically-confirmed “ground truth” values. Next, paired-end simulated datasets with 100 bp read length and simulated sequencing error were generated at 25, 50 and 75 million reads per library, modeled after the same mouse RNA-seq data using polyester (25), which additionally generated ground truth TPM values for each library. Mapping of the simulated reads to GRCm38.p6 (26) was performed with STAR v2.7.8a (27). Ground truth PSI and dPSI values were calculated from the TPM values with a custom Python script that leverages the event-transcript associations in the junctionCounts IOE file using the following equation:

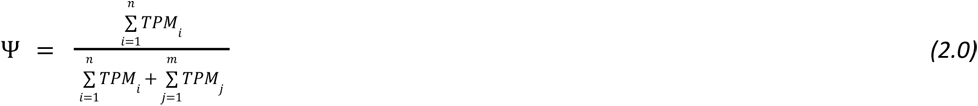

where transcripts *i* in the numerator are those consistent with the included form of the event, and transcripts *j* in the denominator are those consistent with the excluded form of the event.

Events were defined by junctionCounts from the GENCODE M20 (GRCm38.p6) basic gene annotation. Events were defined with splice graphs built by MAJIQ and splAdder in similar fashion; taking the same gene annotation in addition to BAM files from the simulated reads. Event PSI values were measured from the BAM files using each tool’s quantification programs. Each tool’s event naming conventions and definitions were matched to the conventions used by junctionCounts such that the included and excluded forms were consistent in the event definitions, and only events with exact coordinate matches to those in the simulated datasets were considered. Performance on the simulated data was assessed in terms of sensitivity (TPR), false discovery rate (FDR) and quantification error (MARD), calculated accordingly:

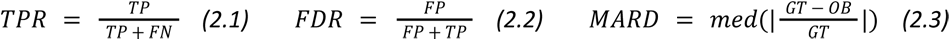

where the true positive rate (*TPR*) is the number of true positives (*TP*) divided by the sum of *TP* and false negatives (*FN*). False discovery rate (*FDR*) is the number of false positives (*FP*) divided by the sum of *FP* and true positives (*TP*), and median absolute relative difference (*MARD*) is the median of the absolute values of ground truth (*GT*) values minus observed (*OB*) values, divided by *GT*.

To make the summary performance comparisons fair, Pearson correlation coefficient was calculated between ground truth PSI/dPSI values and each tool’s observed values for the subset of events that were defined by all three tools. Then to comprehensively assess sensitivity, false discovery rate and quantification error, each tool’s set of events matching events in the simulated data were considered individually. The same performance metrics were also measured for each event type specifically.

### cdsInsertion and findSwitchEvents to couple alternative splicing events to potential functional consequences

cdsInsertion translates transcripts *in silico* from user-provided overlapping start codons and determines resulting transcript characteristics including: UTR lengths, putative protein sequences, the presence of PTCs, PTC distances from downstream splice sites and more. For a given codon and transcript, cdsInsertion first checks whether the start codon’s first position overlaps the genomic coordinates of a transcript’s exons. If it does, the start codon sequence is checked in the spliced transcript’s sequence. Currently, only AUG initiation is supported. If the start codon sequence is AUG, cdsInsertion translates the spliced transcript *in silico* by looking for an in-frame downstream stop codon, which can be: UAA, UGA or UAG. If a downstream stop codon is found, the resulting CDS is recorded along with associated information such as CDS length, coding sequence, PTC presence and PTC distance. If more than one CDS is found within the transcript, additional CDS features are associated to the given transcript as distinct CDS features. If no in-frame downstream stop codon is found, the transcript is recorded as a possible NSD substrate. cdsInsertion outputs a table with summary information about each transcript and a separate GTF file for potential non-PTC, PTC, and non-stop transcript-CDS combinations. The GTF file contains separate transcript records for every CDS-transcript combination. Optionally, cdsInsertion can additionally output bigGenePred files which enable codon visualization on the UCSC Genome Browser. cdsInsertion further outputs a pickled Python dictionary containing all of the aforementioned information associated with each transcript.

Its partner utility, findSwitchEvents, takes an IOE file; a format originally introduced by SUPPA (28), which is generated by junctionCounts to associate events with transcripts, and the pickled Python dictionary containing transcript CDS information and associated details from cdsInsertion. With this information, it evaluates whether isoforms with a specific property (NMD, NSD or CDS) are exclusive to one form of an alternative event. Switch events are AS events that meet this condition, meaning that one (either the included or excluded) form is coupled with a switch to PTC-containing, non-stop or noncoding isoforms within a gene. These tools were evaluated using a previously published dataset in which treatment with emetine, a translation elongation inhibitor, was reported to increase the abundance of NMD substrates in HEK 293 T cells (29). Indeed, cdsInsertion and findSwitchEvents identified 636 potential NMD substrates with significant changes in splicing (|dPSI| ≥ 0.1 and Q-value ≤ 0.05) upon emetine treatment, out of which 472 (74.2%) exhibited the expected dPSI directionality (Supplementary Figure 1).

### Analysis of interspecies and temporal alternative splicing dynamics during neuronal differentiation in primate PSCs

We analyzed RNA-seq data from human and rhesus macaque embryonic stem cells (ESCs) as well as chimpanzee and orangutan induced pluripotent stem cells (iPSCs) (30). Field et al. induced differentiation of the stem cells to cortical neurospheres to model prenatal brain development. Duplicate RNA-seq libraries from each time point (zero, one, two, three, four and five weeks of neuronal differentiation) were downloaded as compressed FASTQ files from SRA, deduplicated and mapped with STAR v2.7.8a to the appropriate genome: GRCh38 (https://www.ncbi.nlm.nih.gov/grc), panTro4 (31), ponAbe2 (32) and rheMac8 (33) for human, chimpanzee, orangutan, and rhesus macaque respectively. The GENCODE v27 (34) basic gene annotation was used as a basis for CAT (35) to generate gene annotations of similar complexity for all species. To reduce the complexity of the input transcriptomes, only basic transcripts were retained. Following this filtration, these annotations were used as input along with the mapped RNA-seq reads for StringTie v1.3.6 (13) to assemble unannotated transcripts. Using the StringTie merge command, comprehensive gene annotations were produced for each species.

To identify orthologous AS events, the whole genome sequences of human (GRCh38), chimpanzee (PanTro4), orangutan (ponAbe2), and rhesus macaque (rheMac8) were mapped to one another using minimap2 v2.11-r797 (36) with parameters *--cs* and *-asm20*. The resulting mappings were used to lift the coordinates of alternative event exons to other species with a modified version (altered such that the input BED file coordinates are semicolon-delimited rather than underscore-delimited in the name field of the output BED file) of the minimap partner utility paftools. Events were reassembled from the lifted coordinates of component exons, and checked for exon count and event type-concordance with the original event. Lifted events passing these checks were proposed as putative orthologs, and then checked against events natively identified in the target species to identify orthologous relationships. Non-one-to-one relationships were removed from consideration.

Temporal (time points 1-5 weeks of neuronal differentiation versus t_0_) and interspecies (paired time points compared across species) AS analyses were performed using junctionCounts. Events with |dPSI| ≥ 0.1 and Q-value ≤ 0.05 across conditions were considered significantly different. Events exhibiting significant splicing differences in at least one temporal comparison for each species were clustered by their temporal PSI trajectories with CLARA (37) using euclidean distance and with 500 iterations. Conservation of primate temporal splicing dynamics was assessed based on concordance with human temporal splicing dynamics with regard to PSI measurements. Genes with mean |dPSI| ≤ 0.1 for events in chimpanzee, orangutan and rhesus macaque relative to human in pairwise comparisons at each time point were categorized as genes with conserved splicing, while those with mean |dPSI| ≥ 0.3 were categorized as genes with non-conserved splicing. Gene ontology analyses for genes in the temporal event clusters and conserved and non-conserved splicing gene sets was done with Metascape (38).

Functional splicing analyses included assessment of switch events and exonic features. Switch events, which we define as events in which all transcripts consistent with one alternative form contain a particular feature while all transcripts consistent with the other form do not, were identified using cdsInsertion and findSwitchEvents (https://github.com/ajw2329/cds_insertion). In NMD switch events one form, but not the other, introduces a premature termination codon (in-frame stop codon ≥ 55 nt upstream of the final exon-exon junction when translated *in silico* from any overlapping consensus coding sequence start codon). In NSD switch events one form, but not the other, results in transcripts lacking an in-frame stop codon. In coding (to noncoding) switch events one form, but not the other, results in transcripts lacking a coding sequence. Exon ontology analysis was performed with Exon Ontology (39) using the included form coordinates of alternative exons as the test list, and the excluded form coordinates as the background.

### Data sources

RT-PCR data matching Zhang et al. (2014) (24) was derived from Vaquero-Garcia et al. (2016) (22) who replicated the experiments in Zhang et al. (2014). Simulated RNA-seq data was modeled on the mouse cerebellum and liver RNA-seq data from Zhang et al. (2014) (24). Published RNA-seq data from emetine-treated and control HEK 293 T cells was taken from Martinez-Nunez et al. (2017) (29). Published RNA-seq data from human and rhesus macaque ESCs as well as chimpanzee and orangutan iPSCs was from Field et al. (2019) (30).

## RESULTS

### junctionCounts benchmarks comparably and identifies a superset of events and event types relative to other tools

We evaluated junctionCounts by comparing its performance on real and simulated data with that of two established AS analysis tools: MAJIQ (22) and splAdder (23). Each publicly available splicing tool identifies and quantifies its own limited set of event types, usually not extending beyond canonical event types and often limited to non-terminal events. splAdder is representative of a tool that only considers non-terminal events and whose definitions correspond to: SE, MS, A3, A5, MX and RI event types in junctionCounts. Additionally, splAdder quantifies events by RNA-seq read coverage at exons and introns rather than by junction reads. MAJIQ identifies relatively numerous event types, the ones matching junctionCounts event types being: SE, MS, A3, A5, MX, AF, AL and RI (with several others that aren’t directly analogous). Similarly to junctionCounts, MAJIQ quantifies these events by junction reads from RNA-seq data, albeit with an approach that is different from junctionCounts. We determined that this was a suitable set of tools for benchmarking junctionCounts as each one has its own merits, identifies events similarly enough to be directly compared across tools and applies distinct quantification approaches.

We first identified alternative events in mouse cerebellum and liver RNA-seq data from Zhang et al. (2014) (24) using junctionCounts, MAJIQ and splAdder. We matched events between the tools based on identical exon-exon and exon-intron junctions in both the included and excluded forms. We initially tested each tool’s PSI and dPSI quantification accuracy on 50 SE events for which biologically confirmed “ground truth” values were generated in RT-PCR data by Vaquero-Garcia et al. (2016) (22), matching the Zhang et al. (2014) (24) RNA-seq dataset. Linearity of observed versus ground truth PSI and dPSI values was assessed by Pearson correlation coefficient (*r*), revealing strong and comparable quantification accuracy by each tool (Figure 1A). Next, we generated simulated datasets modeled on the aforementioned mouse cerebellum and liver RNA-seq data in order to assess each tool’s performance on a much wider set of events and event types with known PSI values, using polyester (25) and a custom Python script that calculates ground truth event PSIs from ground truth transcript TPMs using the event-transcript associations in the junctionCounts IOE file.

**Figure 1.**
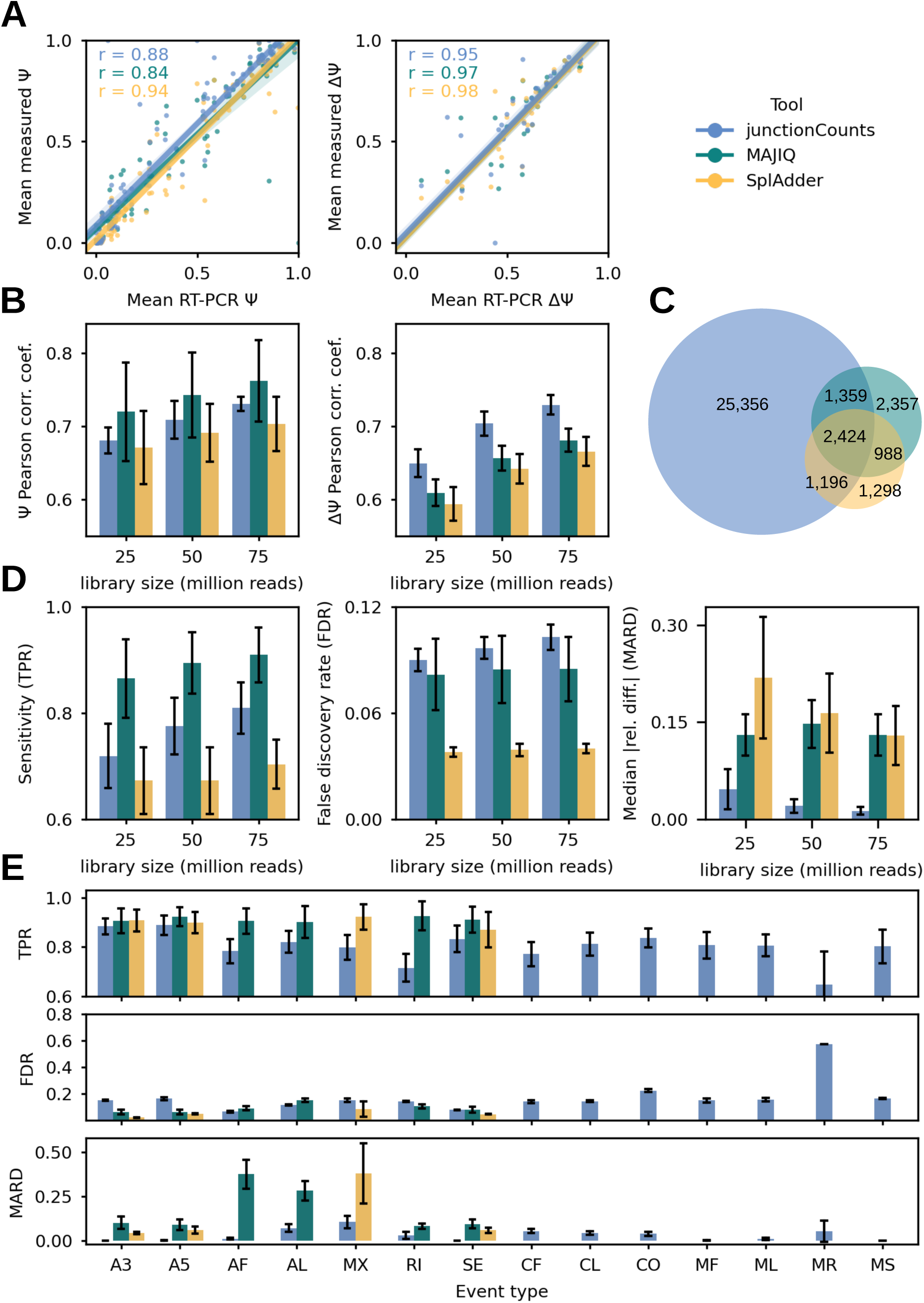
Benchmarking junctionCounts performance relative to similar tools. (A) Comparison of junctionCounts to two established alternative splicing analysis tools: MAJIQ and splAdder. Observed PSI and dPSI values for RNA-seq data, derived from Vaquero-Garcia et al. (2016) (22), compared against RT-PCR-verified values. (B) RNA-seq datasets with known ground truth PSI values, simulated from Zhang et al. (2014) (24) mouse cerebellum and liver RNA-seq data, were generated at 25, 50 and 70 million reads per library in triplicate per tissue type. Observed versus ground truth PSI and dPSI linearity was quantified by Pearson correlation coefficient. (C) Venn diagram of the events identified by each tool in the simulated data. (D) Overall sensitivity (true positive rate), false discovery rate and quantification error (absolute median relative difference) calculated for each tool on the simulated data. (E) The same metrics as in (D) but specific to the 75 million read depth dataset and stratified by event type.

To assess the impact of library depth on event identification and quantification we generated three simulated paired-end RNA-seq datasets of varying depth (25, 50 and 75 million reads per library), each comprising three replicates modeled on mouse cerebellum and three replicates modeled on mouse liver RNA-seq from Zhang et al. (2014) (24). On these datasets, junctionCounts moderately outperformed MAJIQ and splAdder in terms of overall quantification accuracy measured by *r* (Figure 1B). In terms of event identification junctionCounts, MAJIQ and splAdder identified 30K, 7K and 6K events in the 75 million read depth dataset respectively – with MAJIQ and splAdder matching 48-61% of their events with each other and with junctionCounts (Figure 1C). Expectedly, junctionCounts identified a superset of events relative to other programs because it handles complex event types which are uniquely defined and consequently impossible to directly compare to MAJIQ’s non-canonical event types.

Next, we evaluated each tool’s sensitivity (true positive rate or TPR), false discovery rate (FDR) and quantification error (median absolute relative difference or MARD) on the same simulated datasets. junctionCounts had middle-of-the-road sensitivity, slightly inflated FDR and notably low quantification error (Figure 1D). While junctionCounts sensitivity and quantification accuracy scale directly with library read depth, so too does its FDR to a marginal degree. Each tool’s performance improved overall with increased read depth. To appraise event type-specific performance, we calculated the same metrics for each event type on the 75 million read depth dataset (Figure 1E). We found that junctionCounts generally performed well for all event types except for MR events. This is likely because junctionCounts uses the same simple junction read-based approach to quantify all event types, which may confuse ordinary intron contamination for RI or MR events. Another source of bias is that the simulated datasets were modeled on mouse RNA-seq data which contained less than 10 bona fide MR events. To correct for potential false positive RI/MR event calls within a condition, we recommend users filter RI/MR events specifically for mean(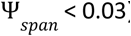) and mean(included junction counts) > 15 across replicates.

Differential splicing analysis across conditions largely eliminates this problem. Taken as a whole, junctionCounts performs on par with similar tools while enabling the identification and quantification of complex event types that are supported by RNA-seq data but are not captured by most available tools.

### Characterizing temporal and species-specific alternative splicing dynamics during primate neuronal differentiation

After establishing that junctionCounts competently characterizes AS in simulated data, we next wanted to examine its utility on real data. To that end, we analyzed a primate neuronal differentiation RNA-seq dataset comprising human and rhesus macaque ESCs as well as chimpanzee and orangutan iPSCs from Field et al. (2019) (30) (Figure 2A). We hypothesized that because the four primates share 90-99% genome sequence conservation (40), junctionCounts should identify a substantial number of orthologous AS events across the four primates (41). We further expected to observe substantial species-specific splicing dynamics during neuronal differentiation as previous interprimate studies have reported (41).

**Figure 2.**
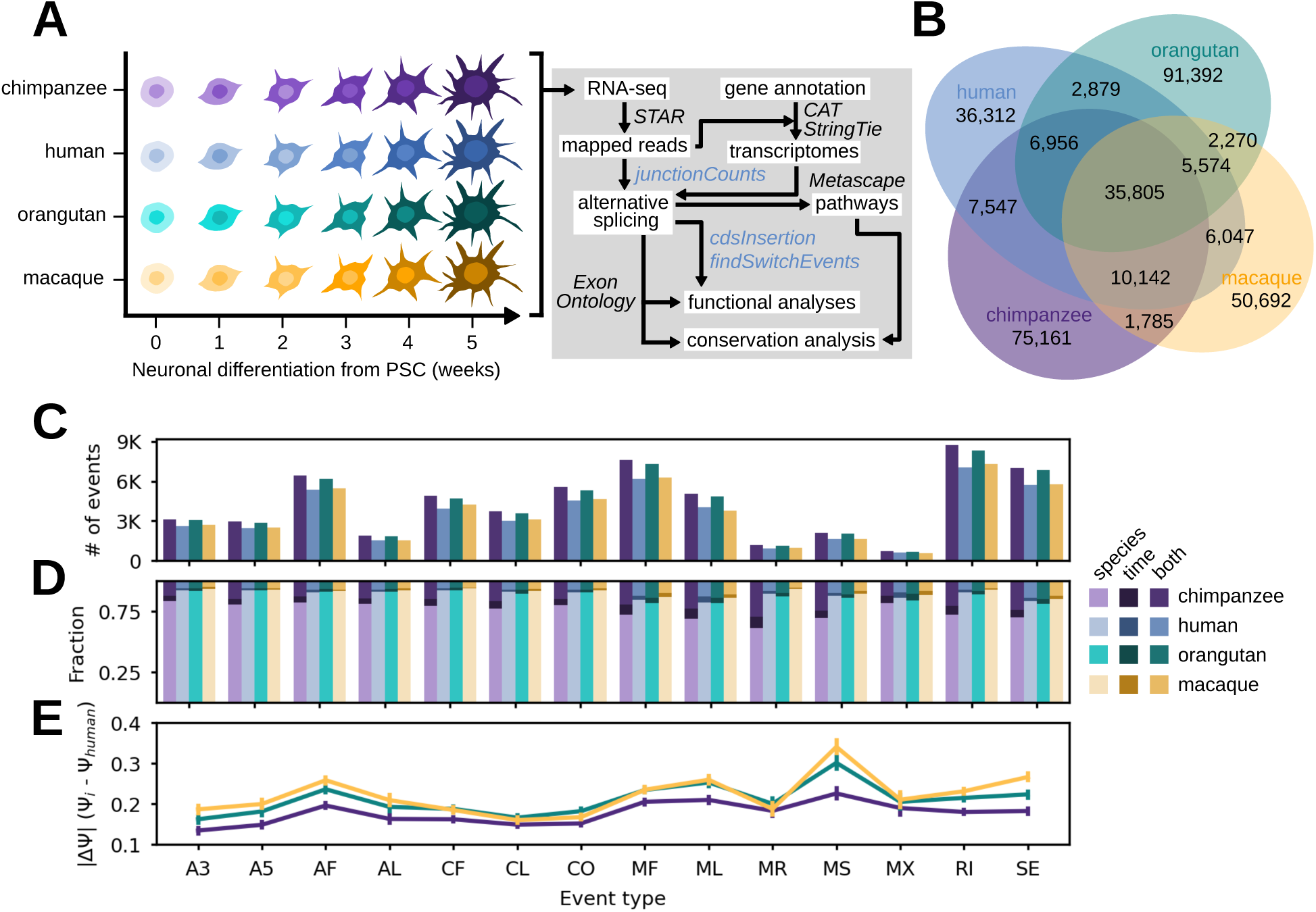
Application of junctionCounts to a primate neuronal differentiation time course experiment. (A) Schematic of five-week chimpanzee, human, orangutan and rhesus macaque neuronal differentiation from pluripotent stem cells and subsequent RNA-seq analysis workflow. (B) Venn diagram of the events identified by junctionCounts in each primate transcriptome. (C) The total number of significant events (|dPSI| ≥ 0.1 and Q-value ≤ 0.05 in at least 1 temporal or interspecies comparison) by event type for each species. (D) The fraction of events that were significantly different across species, time or both factors. (E) Evaluation of conserved splicing by event type, measured by |dPSI| against human PSI values.

We used CAT (35) on the GENCODE v27 (34) basic gene annotation to generate gene annotations of similar complexity for all species. We then used StringTie v1.3.6 (13) on the resultant gene annotations along with the mapped RNA-seq reads to assemble unannotated transcripts. Thus we produced comprehensive gene annotations for each species. Using junctionCounts, we identified approximately 143K, 111K, 151K and 113K possible events in the chimpanzee, human, orangutan and rhesus macaque gene annotations respectively. And to identify orthologous AS events, we performed pairwise mapping of the whole genome sequences of human (GRCh38), chimpanzee (PanTro4), orangutan (ponAbe2), and rhesus macaque (rheMac8) using minimap2 (36). Using the mappings, we lifted the coordinates of alternative event exons to other species using paftools. We then reassembled events from the lifted coordinates of component exons, assessed exon count and event type-concordance with the original events and checked these against events identified in the target species to establish orthologous relationships for which only one-to-one relationships were considered.

In all pairwise interspecies event set comparisons, at least 40% of events were not species-specific, with over 35K orthologous events common to all four primates (Figure 2B). We next quantified these events with junctionCounts which uses junction reads from the mapped RNA-seq data, after which we performed event-level – statistically tested with DEXSeq (21) – pairwise temporal comparisons (week_i_ versus week_0_ of neuronal differentiation) and interspecies comparisons (between corresponding time points; week_i_ versus week_i_) with duplicates per condition. We identified 61K, 50K, 59K and 51K events that were significantly differentially spliced (|dPSI| ≥ 0.1 and Q-value ≤ 0.05) in at least one temporal or interspecies comparison in chimpanzee, human, orangutan and rhesus macaque respectively. We observed that the majority of splicing changes were in interspecies comparisons (Figure 2D), with RI, MF, SE and AF constituting the most commonly differentially spliced event types (Figure 2C). Intriguingly, when we compared orthologous event PSI values by event type between each primate and human across corresponding time points – as a proxy for conservation of splicing dynamics – we found that complex event types (CF, CL and CO) displayed the closest central tendency of PSI values to those of human cells (Figure 2E). This finding may lend credence to the value of characterizing complex event types and their involvement in primate neuronal differentiation.

### junctionCounts uncovers novel splicing dynamics in genes relevant to neuronal differentiation and function

Among the 17K significant events (|dPSI| ≥ 0.1 and Q-value ≤ 0.05 in at least one comparison) that were orthologous in all four primates (Figure 3A), we hypothesized that junctionCounts would both recapitulate previously reported splicing phenomena and identify novel events in genes involved in neuronal differentiation and function. Here, we highlight several such findings. Amphiphysin 1 (AMPH) and Amphiphysin 2 (BIN1) are both enriched in the mammalian brain and participate in synaptic vesicle endocytosis (42, 43). Splice variants of BIN1 have been reported in the brain as well as other tissue types (43), but the implications of AS in AMPH1 remain unexplored. We report a SE event involving exon 17 of AMPH1 (Figure 3B), which is the only scenario of AS that affects the CDS among AMPH1 isoforms annotated in GENCODE V44. According to Exon Ontology (39), AMPH1 exon 17 encodes an intrinsically unstructured polypeptide region which contains an O-phospho-L-serine modification site. This exon is increasingly spliced in over the time course with species-specific trajectories and magnitudes, possibly indicating a functional role for AMPH1 exon 17 inclusion in neurons.

**Figure 3.**
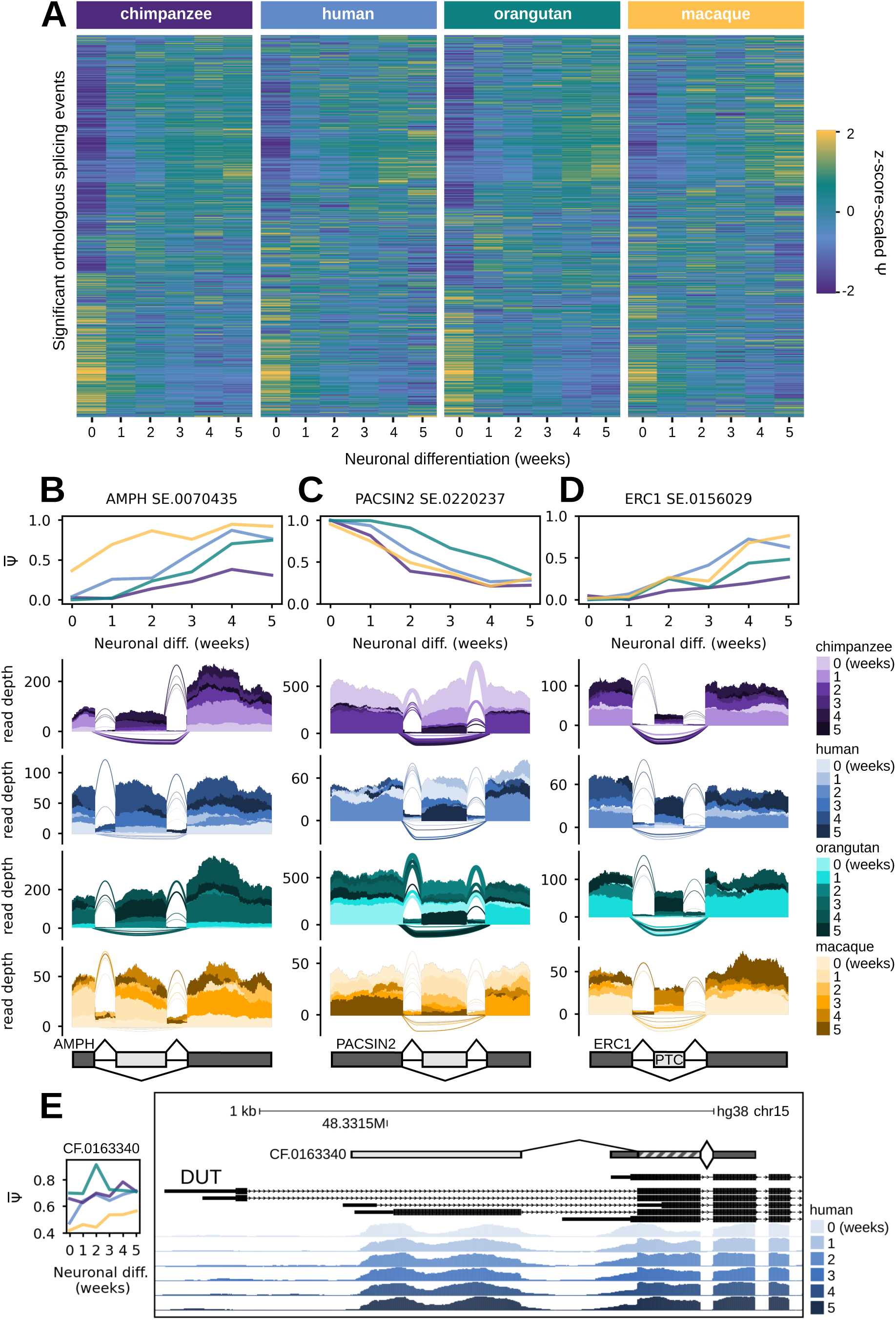
junctionCounts uncovers conserved and species-specific temporal splicing patterns among orthologous splicing events across the four primates. (A) Heatmap of Z-score-scaled PSI values for significant orthologous splicing events (|dPSI| ≥ 0.1 and Q-value ≤ 0.05 in at least 1 temporal or interspecies comparison) with each row corresponding to the same event across all four primates. (B) Mean PSI trajectories and RNA-seq coverage at a skipped exon event in AMPH with species-specific temporal splicing patterns across chimpanzee, human and rhesus macaque. (C) Mean PSI trajectories and RNA-seq coverage at a skipped exon event in PACSIN2 with a conserved temporal splicing pattern. (D) Mean PSI trajectories and RNA-seq coverage at an PTC-containing skipped exon event in ERC1 with a conserved temporal splicing pattern. (E) UCSC Genome Browser snapshot of human read support at a complex first exon event in DUT, which measures the inclusion of one of several distal alternative first exons and its subsequent second exon versus the proximal first exon which overlaps with the alternative second exon. The subpanel to the left shows the mean PSI of the included form at each time point of neuronal differentiation in each primate. In panels (B), (C) and (D) the included form of the alternative event contains both the dark and light gray components, while the excluded form only contains the dark gray components. In panel (E), the same is true except the included form does not contain the dark gray fragment at the 5’ end of the central exon. In panels (B), (C), (D) and (E), the upright and inverted arches represent junction read coverage for the included and excluded form respectively.

Protein kinase C and casein kinase II substrate in neurons 2 (PACSIN2) is the only known member of the PACSINs whose expression isn’t cell type-specific, in humans. All three PACSINs have been reported to play a role in trafficking AMPA receptors in and out of synapses, which is a crucial factor in important neuronal processes including synaptic transmission and plasticity (44, 45). We identified a SE event involving exon 9 of PACSIN2 for which the included form uniformly decreases from the dominant to the minor form over the course of neuronal differentiation (Figure 3C). Similarly to the aforementioned SE event in AMPH1, this SE event is the only CDS-altering event among PACSIN2 isoforms annotated in GENCODE V44. According to the GTEx V8 RNA-Seq Read Coverage by Tissue track on the UCSC Genome Browser (46), PACSIN2 exon 9 inclusion is dominant in all non-neuronal tissue types, while the excluded form is dominant in 9 out of 14 neuronal tissue types. These observations make a compelling case for neuron-specific AS of PACSIN2, resulting in the preferential exclusion of exon 9 in several neuronal cell types.

We discovered a conserved ERC1 PTC-inducing SE event involving exon 18 in isoform ENST00000355446.9 (in GENCODE V44), that to our knowledge has not been previously reported by other groups (Figure 3D). Inclusion of this exon may produce an NMD substrate, but could potentially yield a functional protein isoform at least 30 residues shorter at the C-terminus relative to isoforms consistent with the excluded form. ERC1 has been described to undergo neuron-specific AS and is implicated in important functions including neurotransmitter release and neuronal differentiation (47, 48). Over the five week course of neuronal differentiation, the PTC-inducing SE event follows a consistent pattern of becoming increasingly spliced in across the four primates. Interestingly, AS at the C-terminus of Erc1 in rats was shown to generate two isoforms: Erc1a and Erc1b. The latter of which is the brain-specific, shorter isoform that alone can bind to presynaptic active zone proteins, called RIMs, that regulate neurotransmitter release (49). Taken together, these observations suggest a potential functional role for the inclusion of the ERC1 poison exon in differentiating neurons.

Deoxyuridine 5′-triphosphate nucleotidohydrolase (DUT) is an important enzyme involved in genome integrity maintenance that prevents uracil misincorporation into DNA. DUT expression has been shown, through knockout studies, to be essential to embryonic development and especially to later stages of differentiation in mice (50). We identified a CF event in DUT (Figure 3E), in which the included form corresponds to the DUT-M isoform and the excluded form corresponds to the DUT-N isoform (51). The DUT-M isoform localizes to mitochondria via a mitochondrial targeting presequence located in the first exon consistent with the included form of the CF event and is expressed constitutively. The DUT-N isoform localizes to the nucleus and its expression is induced during the G_0_ to S phase transition. Exit from the cell cycle into G_0_ phase triggers DUT-N protein degradation. Thus, DUT-N isoform expression is tightly linked to nuclear DNA replication (51). We observed that the CF event is increasingly spliced in – meaning DUT-M isoform expression gradually eclipses DUT-N isoform expression over the time course – which is to be expected as the primate pluripotent stem cells (PSCs) progressively commit to neuronal cell fates with decreasing cell cycle activity (52). These findings demonstrate that junctionCounts can handily uncover novel splicing phenomena.

### Temporal regulation of alternative splicing directs the transition from pluripotent to neuronal cell fate

High levels of AS and cell type-specific isoform expression are observed in neurons and during neuronal differentiation (53, 54). We postulated that genes exhibiting dynamic temporal splicing would be enriched for neuronal biological pathways. Taking the subset of significant AS events (|dPSI| ≥ 0.1 and Q-value ≤ 0.05 in at least one temporal comparison), we generated the four most distinct clusters of events based on the Euclidean distance of their temporal PSI trajectories using CLARA (37) for each species (Figure 4A). Additionally, we identified subsets of genes with conserved (mean |dPSI| ≤ 0.1 for all events per gene) and nonconserved (mean |dPSI| ≥ 0.3 for all events per gene) splicing patterns in chimpanzee, orangutan and rhesus macaque relative to human in pairwise comparisons at each time point (Figure 4B). We then used Metascape (38) to identify enriched biological pathways in the sets of genes from each cluster of temporally regulated events (Figure 4C) and for the conserved and nonconserved splicing gene sets (Supplementary Figure 2).

**Figure 4.**
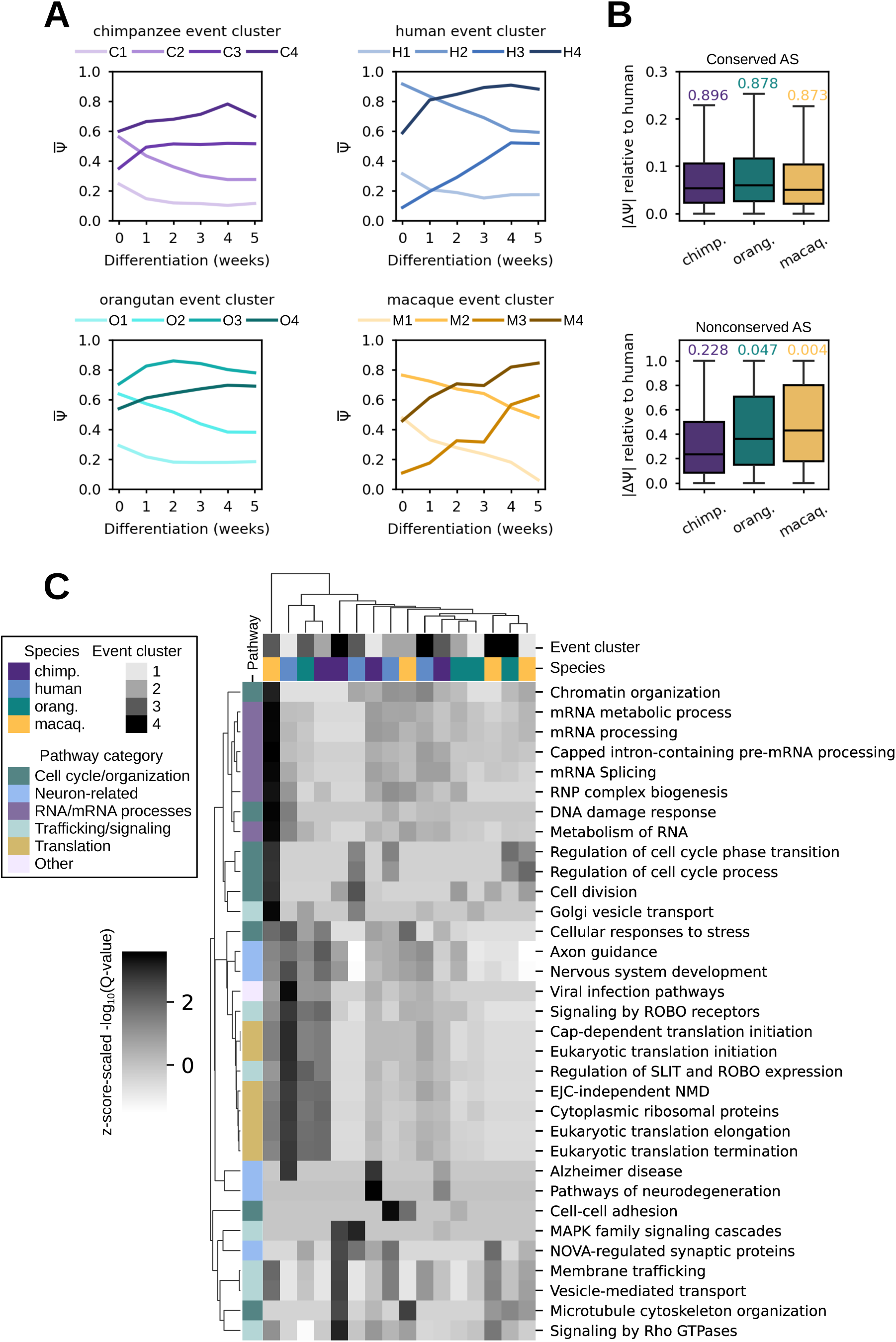
Gene ontology analyses for grouped event sets by temporal PSI trajectories. (A) Four gene-level clusters derived with CLARA from temporal expression trajectories for each species. (B) Genes with mean |dPSI| ≤ 0.1 for events in chimpanzee, orangutan and rhesus macaque relative to human were categorized as genes with conserved splicing. Boxplots showing the distribution of each primate’s per-gene mean |dPSI| relative to human (upper subpanel) with Pearson correlation coefficient of species-specific PSI values against human PSI values above them. The same for genes with nonconserved splicing based on mean |dPSI| ≥ 0.3 (lower subpanel). (C) Heatmap of Z-score-scaled-log_10_(Q-value) of Metascape (38) pathway enrichment in temporal event clusters.

Four similar but distinct event clusters were identified in each primate. Across the primates, cluster 1 (chimpanzee, human, orangutan and rhesus macaque corresponding to C1, H1, O1 and M1 respectively) generally represents alternative features (exons, introns, splice sites, etc.) whose inclusion is marginal in PSCs and declines over the course of differentiation. Cluster 2 (C2, H2, O2 and M2) represents alternative features whose inclusion is dominant in PSCs and declines during differentiation. Pathways similarly enriched in clusters H1, M1 and C2 indicate the preferred exclusion of particular alternative features in the mature splicing program of genes related to: translation, NMD, axon guidance and nervous system development. Cluster 3 (C3, H3, O3 and M3) generally contains alternative features whose inclusion is marginal in PSCs and increases during differentiation, while cluster 4 (C4, H4, O4 and M4) contains dominantly included alternative features that further increase until peaking at week 4 during differentiation. Clusters H3 and O3 indicate increasing inclusion of alternative features in the mature splicing program of genes related to: NOVA-regulated synaptic proteins, axon guidance and nervous system development. Cluster C3 indicates a slight increase in alternative feature inclusion in genes related to mRNA processing and translation. The conserved splicing event set was enriched for genes in critical pathways including cell cycle processes, signaling, AS, and interestingly, in neurodegeneration pathways. The subset of complex conserved splicing events (CF, CL and CO) was enriched for nearly all the same pathways, revealing the prevalence of complex events in important pathways (Supplementary Figure 2). For example, we identified a conserved CO event in NCKAP1, which is involved in Rho GTPase signaling, and a conserved CL event in QKI, which is involved in pre-mRNA processing and AS (Supplementary Figure 3). Taken together, these results highlight the intricate temporal regulation of splicing as PSCs develop into neuronal cells and shed light on the biological relevance of species-specific and conserved splicing dynamics.

### Emergent alternative features underlie many instances of species-specific alternative splicing

Because we observed that some events had miniscule or zero PSI values in particular species, we hypothesized that a subset of the aforementioned nonconserved event set represents events that sufficiently map (sequence divergence ≤ 20%) pairwise between all four primate genomes but contain alternative features that are only used (included) by specific primates despite the apparent presence of splice site and branch site sequences. We call features used by specific species emergent alternative features, potentially indicating exonization events. Indeed, we identified 3753 events, in 1922 genes, exhibiting significant temporal regulation (|dPSI| ≥ 0.1 and Q-value ≤ 0.05 in at least one temporal comparison) while having a min(PSI) ≥ 0.05 in only a subset of the four primates (Figure 5A). Of these events, the most prominent event types were SE, MF, AF and ML (Figure 5B). Protection of Telomeres 1 (POT1) is an example of a rhesus macaque-specific SE event in the 5’UTR (Figure 5C). This SE event is likely an instance of species-specific differences in exon induction related to neuronal differentiation, as the alternative exon is annotated in GENCODE V44 and exhibits cell type-specific expression in a number of human neuronal tissues according to GTEx V8 RNA-Seq Read Coverage by Tissue despite its lack of inclusion in our human samples. Transmembrane Protein 165 (TMEM165) is an example of a human-specific PTC-containing SE event that is potentially the product of Alu exonization (55), as it overlaps an antisense AluJb element (Figure 5D). Furthermore, one piece of evidence that suggests that it may be a bona fide emergent alternative cassette exon is that the rhesus macaque genome sequence has an A→G mutation 3 nt upstream of the 3’SS while the other three primates have a canonical 3’SS dinucleotide (Figure 5E). In short, identification of AS events in orthologous sequences between species may be an effective approach to uncover potential emergent alternative features.

**Figure 5.**
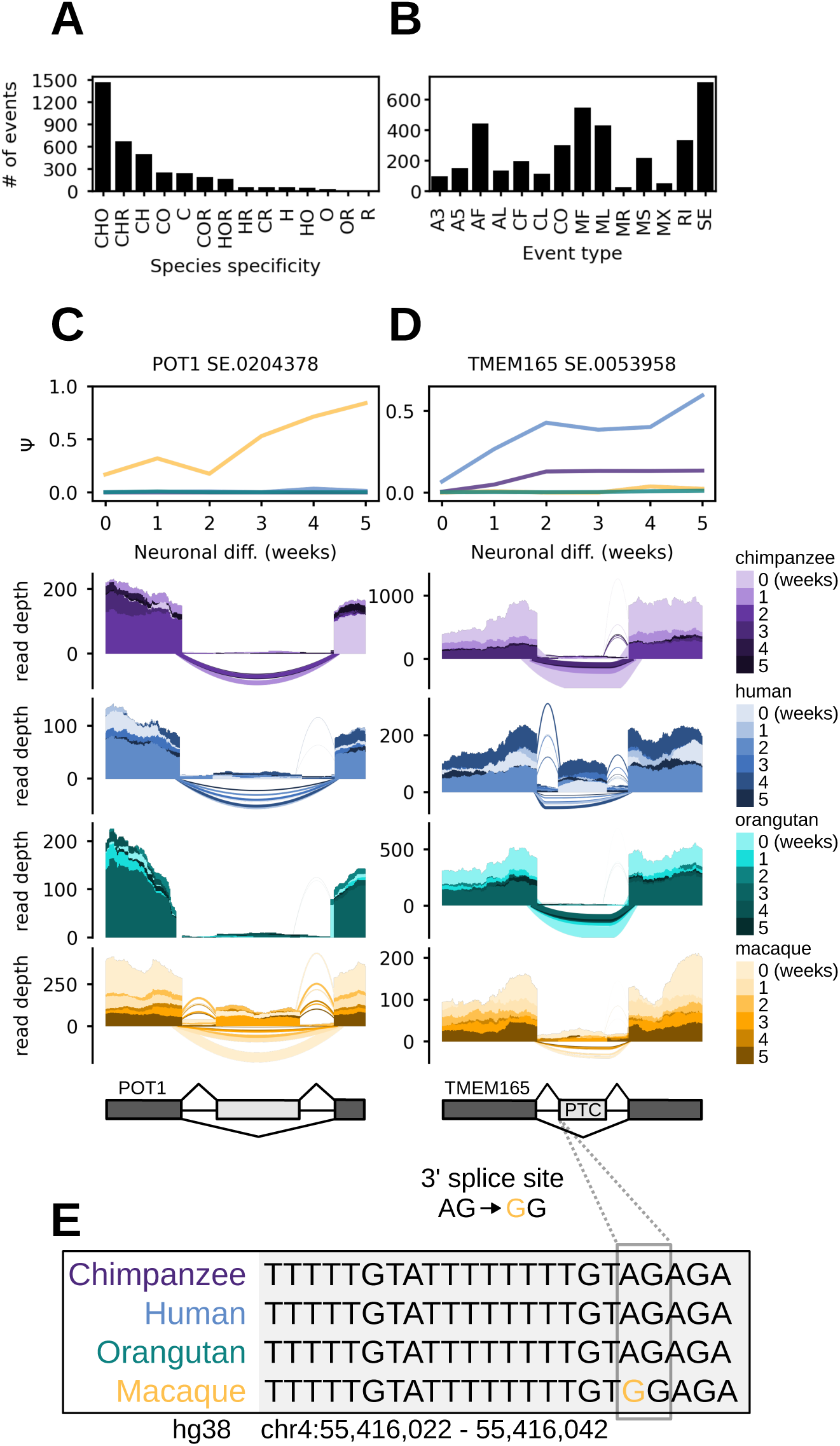
Emergent/species-specific alternative feature usage. (A) Barplot showing the number of species-specific events (min(PSI) ≥ 0.05 in a given species) exhibiting significant temporal regulation (|dPSI| ≥ 0.1 and Q-value ≤ 0.05 in at least 1 temporal comparison). “C”, “H”, “O” and “R” are abbreviations for chimpanzee, human, orangutan and rhesus macaque respectively. Combinations of these abbreviations represent instances of events that meet the min(PSI) threshold in a set of species and are significantly temporally regulated in at least 1 species in the subset. (B) Barplot displaying the same set of species-specific events as in (A) but stratified by event type instead of species. (C) Mean PSI trajectories and RNA-seq coverage at a rhesus macaque-specific skipped exon event in POT1. (D) Mean PSI trajectories and RNA-seq coverage at a human-specific skipped exon event in TMEM165. (E) Macaque-specific point mutation just upstream of the 3’SS of the TMEM165 skipped exon event shown in (D).

### cdsInsertion and findSwitchEvents connect alternative splicing events to potential functional impacts

An enduring problem in the study of AS is the challenging nature of connecting events to functional impacts, whether at the mRNA or protein level. We used cdsInsertion to annotate transcripts with information regarding the lengths of the UTRs and CDS, the presence of potential PTCs and other details gleaned from overlapping annotated start codons. Next, we employed findSwitchEvents to couple transcript-level CDS information to junctionCounts-defined events to identify “switch events”, which are instances in which a particular property is exclusive to transcripts consistent with the included or excluded form. We propose that this approach enables users to connect AS events to functional outcomes, comprehensively profile switch event regulation and to discover novel instances of NMD/NSD.

Among significant events (|dPSI| ≥ 0.1 and Q-value ≤ 0.05 in at least one comparison), we identified hundreds of events predicted to confer NMD, NSD and coding-to-noncoding switches as well as >1600 CDS-altering events in each species (Figure 6A). To investigate the potential structural and functional impacts of CDS-altering events, we mapped event coordinates to protein features with Exon Ontology (39). The five protein feature categories most frequently overlapping alternative exons were: post-translational modification (PTM), structure, binding, localization and catalytic activity (Figure 6B). Interestingly, intrinsically disordered regions (IDRs) were the most highly represented feature (Figure 6C). In agreement with these findings, IDRs have been described as preferred loci for both AS and PTMs (56, 57).

**Figure 6.**
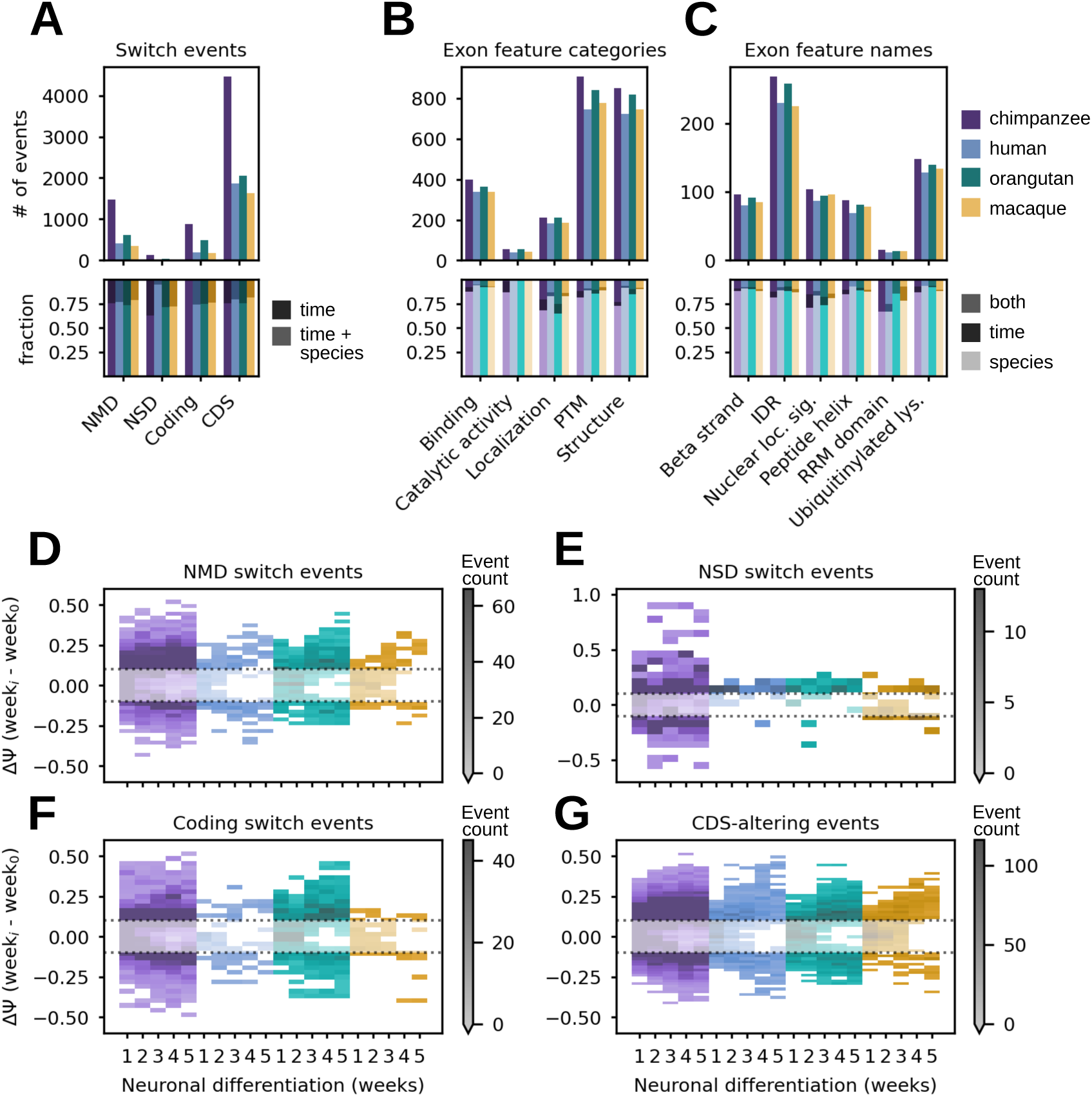
Analysis of splicing events with potential functional impacts to mRNA stability, coding capacity or protein function. (A) Total number of significant splicing events (|dPSI| ≥ 0.1 and Q-value ≤ 0.05 in at least 1 temporal or interspecies comparison) that are predicted to: induce nonsense-mediated decay (NMD), induce non-stop decay (NSD), break the open reading frame (coding switch) or alter the coding sequence (CDS). (B) Exon feature category support for significant event coordinates overlapping exon features in the Exon Ontology database. (C) Specific exon features affected by significant events, indicating potential impacts to structural or functional protein elements. The fraction of events in (A), (B) and (C) that were significantly different across species, time or both factors (lower subpanels). (D) Bivariate histogram showing the distribution of NMD switch event dPSIs over the time course of neuronal differentiation relative to pluripotent stem cells (week 0) in each primate. (E), (F) and (G) show the same for NSD switch, coding switch and CDS-altering events, respectively.

At the mRNA-level, NSD and NMD substrates are expected to be degraded via translation-dependent pathways to prevent the production of potentially harmful truncated proteins (58). However, in certain contexts expression and translational activation of NMD substrates can play important roles in biological functions, including in neuronal differentiation (59). Another possible outcome of AS is the generation of noncoding transcripts from protein coding genes (60). We characterized the regulation of these three phenomena during neuronal differentiation and found that NMD and coding-to-noncoding switch events became progressively more differentially spliced over the course of neuronal differentiation relative to PSCs (Figure 6D-F). NSD switch events were relatively rare, but they were surprisingly overrepresented in chimpanzee relative to the other primates (Figure 6E). Overall, we did not observe a monotonic increase in NMD substrate abundance during neuronal differentiation and we found that NMD/NSD/coding-to-noncoding switch events generally followed species-specific temporal trajectories (Supplementary Figure 4). CDS-altering events were also increasingly differentially spliced over the course of differentiation (Figure 6G), likely owing to the gradual definition of a mature splicing program that requires cell-type specific expression of isoforms with distinct functions (61).

## DISCUSSION

This paper describes our efforts to develop an accurate, rigorous, easy to use and interpret alternative splicing analysis tool capable of identifying and quantifying complex splicing events, and importantly, coupling them to predicted transcript-level functional outcomes. junctionCounts recapitulates both RT-PCR-derived PSI values and ground truth PSI values from simulated data and performs well in our benchmarking experiment against MAJIQ (22) and splAdder (23) (Figure 1). It identifies and accurately quantifies an array of event types including complex event types not captured by most other tools. We note, however, that junctionCounts performed poorly on the MR event type specifically in our benchmarking experiment, likely due to bias within the simulated datasets themselves. Improvement of retained intron event definition and quantification are a priority for future installments. In contrast to MAJIQ and splAdder, junctionCounts identifies events from a gene annotation alone and consequently doesn’t filter events by mapped junction read support. While this eliminates the need to generate new splice graphs/event dictionaries for each individual dataset, it necessitates post-quantification filtering of events with insufficient read coverage. We employ DEXSeq (21) for statistical testing across conditions, which effectively extracts well-supported and significantly regulated splicing events. Contrarily, MAJIQ, splAdder and similar tools use information from mapped RNA-seq data during splice graph generation, so event identification understandably scales directly with library read depth and can include novel splice junctions. Novel splice junction detection can be achieved with junctionCounts by providing a transcriptome assembled from RNA-seq reads using a tool like StringTie (13).

We applied junctionCounts to a primate neuronal differentiation RNA-seq dataset comprising human and rhesus macaque ESCs as well as chimpanzee and orangutan iPSCs from Field et al. (2019) (30). We identified 50-61K significant splicing events (|dPSI| ≥ 0.1 and Q-value ≤ 0.05 in at least one temporal or interspecies comparison) in each species, with 17K significant events that were orthologous across all four primates (Figure 3A). Within these orthologous events, junctionCounts recapitulated previously reported splicing phenomena (51) and identified previously unreported events in several genes relevant to neuronal differentiation (Figure 3B-E). RT-PCR experiments were used to verify some of these events, including SE events in GABBR1 and MYCBP2 in human and macaque cells (Supplementary Figure 5). We additionally clustered events by their temporal splicing dynamics, uncovering distinct event trajectories that capture the tight regulation of splicing during development (Figure 4A). Highly relevant biological pathways were represented in these event clusters, including axon guidance, nervous system development and SLIT/ROBO signaling. Pathways in chromatin organization, mRNA processing and cell cycle processes were also shown to undergo and potentially underlie splicing regulation (Figure 4C).

Within the set of events with nonconserved splicing patterns, we uncovered thousands of events containing alternative features that were used in some primates but not in others, suggesting potential emergent alternative features (Figure 5). Lastly, we used cdsInsertion and findSwitchEvents to connect events to predicted NMD/NSD/coding-to-noncoding switches based on transcript-level CDS properties exclusive to their included or excluded forms. This allowed us to profile temporal NMD/NSD regulation (Figure 6D-E) and to identify potential NMD substrates (Figure 3D and 5D). Altogether, we exhibited the functionality of junctionCounts in a variety of analysis contexts and presented its application to the characterization of splicing in evolution, neuronal differentiation and NMD.

Besides this work, we further note that several of our colleagues have already implemented and published results using a beta version of junctionCounts. These studies include a variety of model systems such as human and non-human primate cell lines, *C. elegans*, and yeast. Suzuki et al. (2022) looked at the effects of KIN17 and PRCC mutations on 5’ and 3’SS usage during development in *C. elegans*. They found both direct and potentially indirect changes in alternative 5’ and 3’SS usage, some of which were related to developmental and population dynamics. They additionally RT-PCR-verified a number of these events to differentiate between embryonic-type splicing and somatic-type splicing (62). Cartwright-Acar et al. (2022) characterized splicing changes in the presence of class II suppressors of uncoordination in an *unc-73(e936)* mutant forward genetic screen in *C. elegans*. They found that the majority of alternative 5’SS usage changes were in introns containing true alternative 5’SS and that suppressors rarely activated novel cryptic alternative 5’SS. They further RT-PCR verified several of the alternative 5’SS and 3’SS events, and finally asserted that the class II suppressors they studied may work at mutually exclusive stages of spliceosome assembly or use different mechanisms to maintain 5’SS identity based on their ability to differentiate between alternative 5’ splicing events that are unique to particular suppressors (63). Draper et al. (2023) quantified events across polyribosome fractions and between primates to assess the conservation of alternative splicing coupled to translational control (ASTC). They identified subsets of alternative events with either conserved or species-specific sedimentation profiles and discovered that alternative exons with conserved sedimentation had higher sequence conservation relative to those with species-specific sedimentation. They additionally tested three ASTC SE events using translational luciferase reporters (64). Hunter et al. (2023) examined the effect of splicing inhibitors on intron splicing efficiency in *S. cerevisiae*. They found that individual introns had distinct sensitivities, including during co-transcriptional splicing, to different splicing inhibitors. Interestingly, they found that yeast sequences including the branch point consensus motif contribute to the differences in sensitivity (65). Osterhoudt et al. (2024) explored changes in 3’SS usage upon SACY-1 perturbation in introns with pairs of 3’ splice sites ≤ 18 nucleotides away from each other. They found that both SACY-1 depletion and a SACY-1 mutation lead to a clear unidirectional increase in proximal alternative 3’SS usage, which they RT-PCR-verified for several events (66). Collectively, our collaborators found junctionCounts easy to implement and show, through these works, its capacity to generate high quality results.

Beyond its flexibility and user-friendliness, junctionCounts stands out as a useful approach because it identifies both canonical and non-canonical alternative events. Many tools are limited to non-terminal and/or relatively rudimentary event types. The few that characterize complex or non-canonical event types are difficult to interpret. junctionCounts utilizes the concept of binary alternative events; identifying clear instances of the inclusion and exclusion of alternative features. This concept is well-established and pervades the splicing literature. It remains popular because binary alternative events can be accurately quantified relative to full-length transcripts, they likely accurately represent (a subset of) transcript structure as compared with full-length transcripts, and they exclude gene segments not relevant to the regulation of the event (i.e. introns and exons outside and distal to the event). The first two reasons will likely decrease in validity as improving long-read sequencing approaches provide more accurate representations of the ground-truth expressed transcriptome. The biggest problem may lie with the third reason, considering that the contribution of factors not necessarily local to an event itself can be important to its regulation (67, 68).

At present, however, focusing on the local site of alternative events affords the opportunity to consider the behavior of hundreds or thousands of similar events and to look for trends in features that may explain their behavior. However, a key weakness of the traditional binary event is the existence of loci in which more than two alternative sub-transcripts overlap and are subject to simultaneous changes in relative abundance. While such non-binary events could be represented as the collection of binary events involving all possible pairs of sub-transcripts, this representation loses information as the regulatory decision is likely to be made in the context of all possibilities. A number of efforts such as MAJIQ (22) and Whippet (17) have attempted to address this issue with several approaches. Nonetheless, junctionCounts presents a step in the right direction by characterizing events that don’t fit into canonical binary event definitions.

junctionCounts furthermore bridges the gap between event-level and transcript-level analysis with regard to the implications of AS events on transcript coding and translational capacity, via cdsInsertion. In an ideal case, studies that intend to consider translation and its implications on a transcriptome-wide scale would include an experimental technique to empirically define CDS regions or start codons. For example, ribosome profiling and approaches like TI-seq (69) can serve as a basis for empirically defining whole CDS or translation start sites respectively. However, as such data are typically unavailable due to the additional cost and complexity of these approaches, tools like cdsInsertion are useful. cdsInsertion fleshes out the putative characteristics of unannotated transcripts by performing *in silico* translation from known overlapping start codons, and thus permits the development of hypotheses to explain properties imparted by AS. Its partner tool, findSwitchEvents, infers alternative event characteristics from those of its constituent transcripts. For instance, it predicts whether or not an event can be considered an NMD switch event. We evaluated those particular predictions using published RNA-seq data profiling changes to NMD substrate abundance upon emetine treatment (29) (Supplementary Figure 1), for which our results support the assertion that findSwitchEvents is capable of predicting NMD switch events and is thus an effective approach for the interpretation of functional alternative event outcomes. Taken together, junctionCounts, cdsInsertion and findSwitchEvents present an easy to install, easy to use and powerful tool for characterizing AS and coupling events to potential functional outcomes.

## DATA AVAILABILITY

Mouse cerebellum and liver RNA-seq data from Zhang et al. (2014) (24) used in benchmarking experiment: publicly available at the NCBI GEO (Accession no.: GSE54652).

RT-PCR data matching Zhang et al. (2014) (24) from Vaquero-Garcia et al. (2016) (22) used in benchmarking experiment: publicly available in their Supplementary Materials Figure 4 source data 1.

RNA-seq data from emetine-treated HEK 293 T cells from Martinez-Nunez et al. (2017) (29) used to validate cdsInsertion and findSwitchEvents NMD predictions: publicly available at the NCBI GEO (Accession no.: GSE89774).

RNA-seq data from human and rhesus macaque ESCs as well as chimpanzee and orangutan iPSCs from Field et al. (2019) (30): publicly available at the NCBI GEO (Accession no.: GSE106245).

The version of junctionCounts and partner utilities used in this study are published on Zenodo (<u>10.5281/zenodo.10674489</u>). junctionCounts and its partner utilities are also available on GitHub: https://github.com/ajw2329/junctionCounts and https://github.com/ajw2329/cds_insertion.

## SUPPLEMENTARY DATA

Supplementary data are available at NAR online.

## ACKNOWLEDGEMENTS

We thank Sol Katzman, Julia Philipp and Jolene Draper for discussion on junctionCounts development and implementation. We thank Timothy Sterne-Weiler for discussion of benchmarking studies and feedback on the manuscript.

---

**Figure S1.**
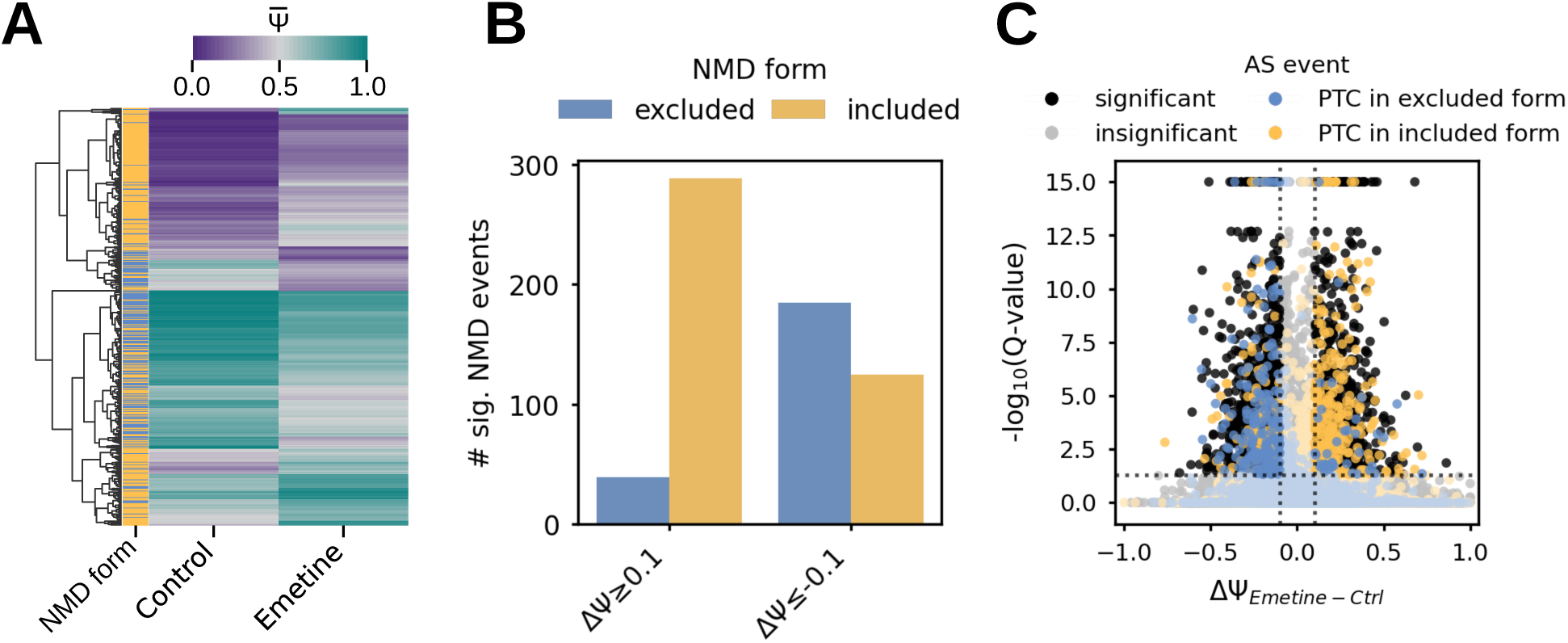
cdsInsertion and findSwitchEvents NMD substrate prediction validation on emetine treatment dataset. Emetine is a translation elongation inhibitor that was reported to increase NMD substrate half-life in HEK 293 T cells (Martinez-Nunez et al. 2017) (29). We applied junctionCounts, cdsInsertion and findSwitchEvents to the RNA-seq data from that study to characterize the splicing of potential NMD substrates based on the predicted presence of a PTC. (A) A heatmap of mean PSI values for predicted NMD events in emetine treatment and control conditions. The NMD form refers to the form confering the PTC: included (yellow) or excluded (blue). (B) A barplot showing the sum of significantly differentially spliced potential NMD substrates exhibiting either a positive or negative dPSI. (C) A volcano plot showing the splicing changes upon emetine treatment, with potential NMD substrates annotated with their associated switch form.

**Figure S2.**
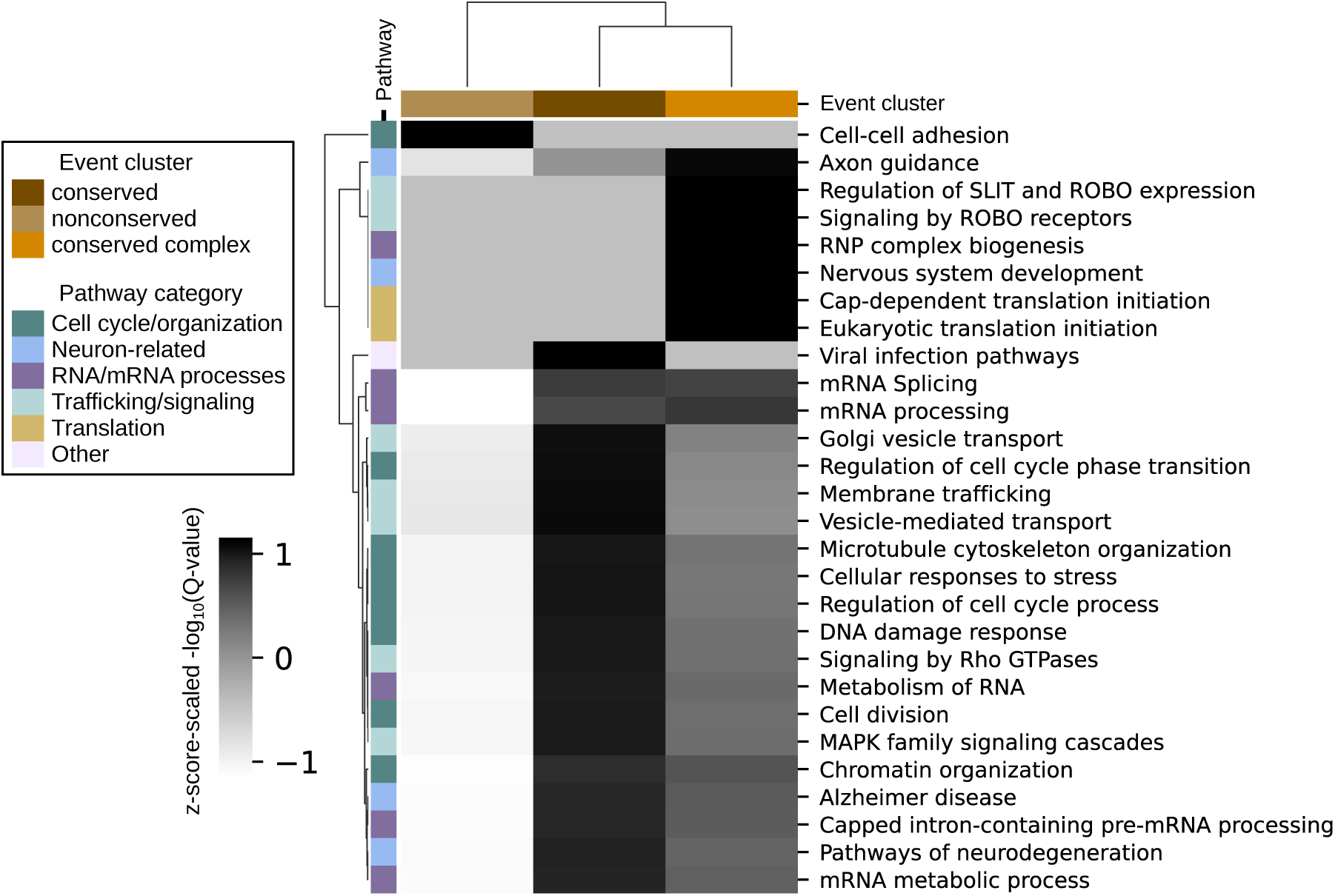
Metascape pathways for conserved and nonconserved event sets. Heatmap of Z-score-scaled-log10(Q-value) of Metascape (38) pathway enrichment in event clusters. “Cons”, “cons_complex” and “noncons” refer to the set of genes with conserved splicing, the subset of genes with conserved complex splicing events, and the set of genes with nonconserved splicing dynamics respectively.

**Figure S3.**
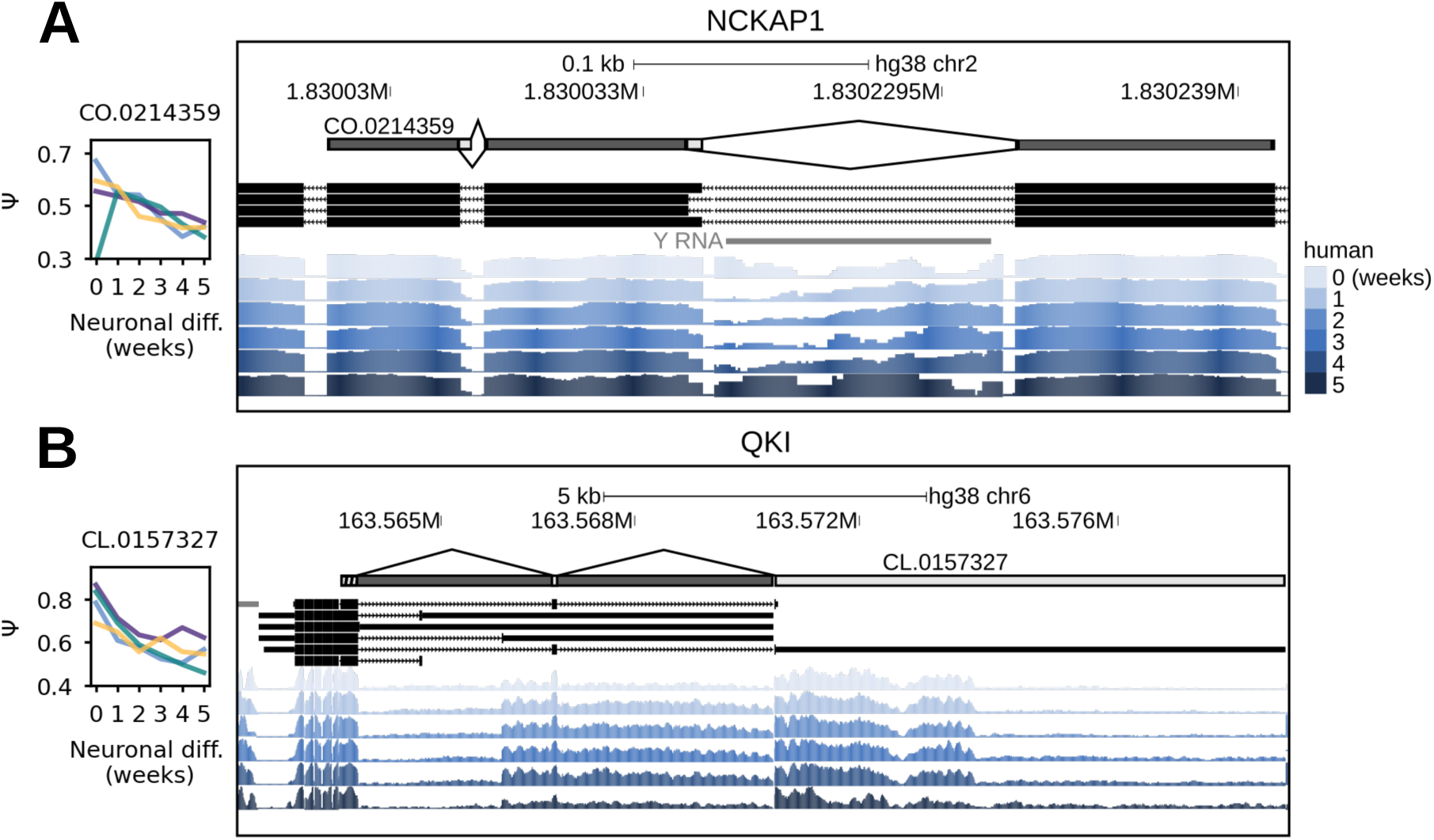
Examples of conserved complex events. UCSC Genome Browser snapshot of human read support at (A) a complex internal event in NCKAP1 and (B) a complex last exon event in QKI. The included form uses the splice junctions above the model and the excluded form uses those below it. PSI trajectory subplots to the left show the event’s inclusion at each time point for chimpanzee, human, orangutan and macaque in purple, blue, green and yellow respectively.

**Figure S4.**
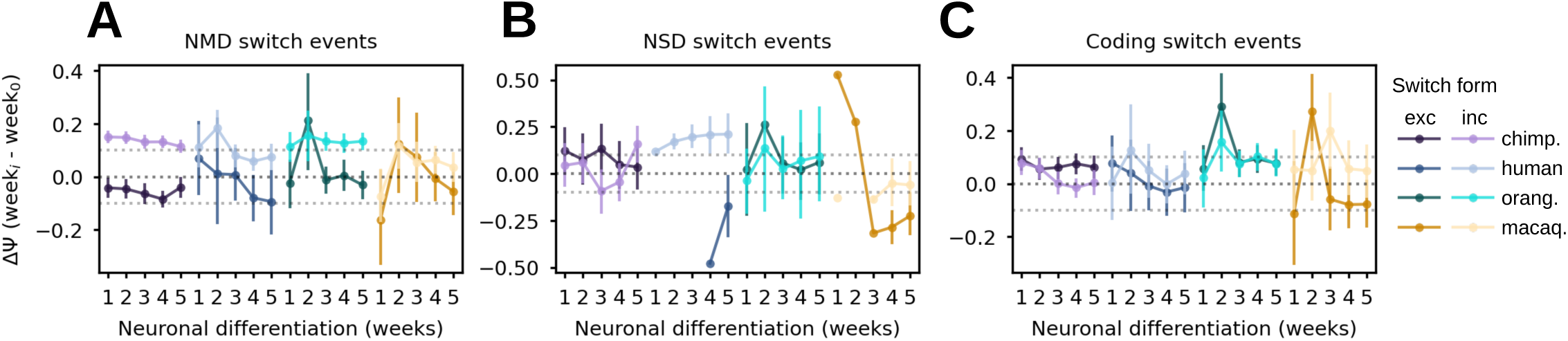
Directionality of switch forms in NMD, NSD and coding-to-noncoding switch events during primate neuronal differentiation. We expect that when the included form of a switch event confers a property (NMD/NSD/noncoding), its increased inclusion (dPSI) signals an increase in its abundance. If the excluded form confers the property, its decreased inclusion signals an increase in its abundance. (A-C) dPSI trajectories of NMD, NSD, coding-to-noncoding switch events respectively relative to week 0 over the time course. The switch form refers to the form confering the property: included (lighter color) or excluded (darker color).

**Figure S5.**
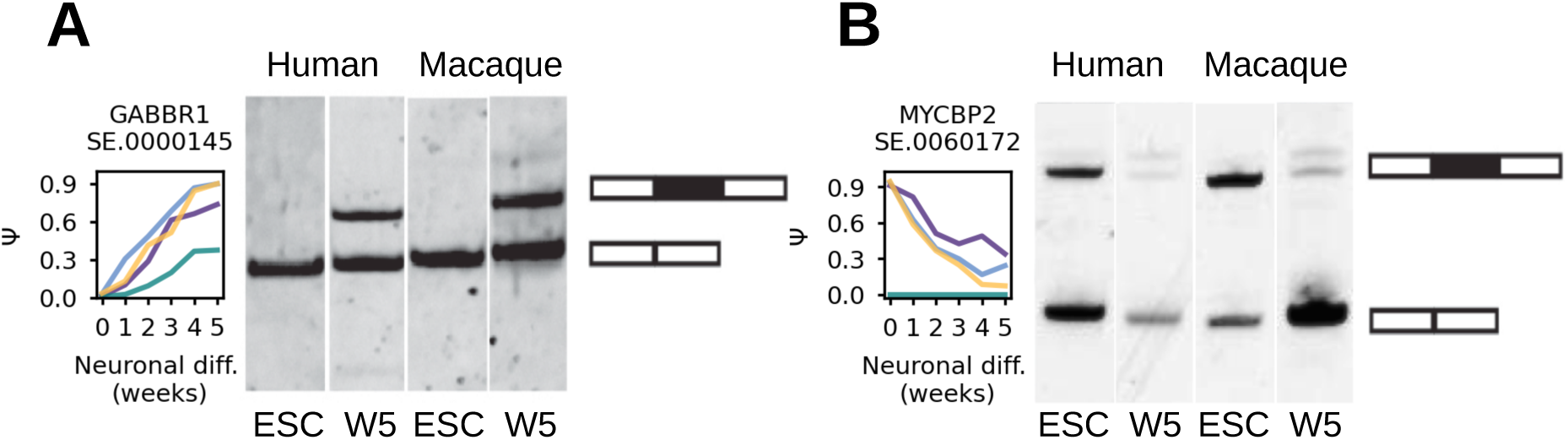
RT-PCR validation of two conserved SE events in human and macaque. (A) An SE event in GABBR1 which is completely excluded in ESC, but gradually included to a similar degree over the course of neuronal differentiation. (B) An SE event in MYCBP2 for which the included and excluded forms are similarly abundant in ESC, but by week 5 the excluded form becomes dominant. These RT-PCR data corroborate our findings by junctionCounts. PSI trajectory subplots to the left show the event’s inclusion at each time point for chimpanzee, human, orangutan and macaque in purple, blue, green and yellow respectively.

